# Inteins in the Loop: A Framework for Engineering Advanced Biomolecular Controllers for Robust Perfect Adaptation

**DOI:** 10.1101/2022.08.05.502923

**Authors:** Stanislav Anastassov, Maurice Filo, Ching-Hsiang Chang, Mustafa Khammash

## Abstract

Homeostasis is one of the cornerstones of life shaped by billions of years of evolution. A notion that is similar to homeostasis, but yet more stringent, is Robust Perfect Adaptation (RPA). A system is endowed with RPA if it is capable of driving a variable of interest to a prescribed level despite the presence of disturbances and uncertainties in the environment. Designing and building biomolecular controllers capable of achieving RPA have been identified as an important task which has immediate implications for various disciplines. Here, we develop systematic theoretical and experimental frameworks for custom-built proteins that exploit split inteins — short amino acid sequences capable of performing protein-splicing reactions — to design, genetically build and analyze a wide class of RPA-achieving integral feedback controllers. We first lay down a theoretical foundation that facilitates the screening of intein-based controller networks for RPA, and then usher an easy-to-use recipe to simplify their, otherwise complex, underlying mathematical models. Furthermore, we genetically engineer and test various controller circuits based on commonly used transcription factors in mammalian cells. We experimentally and theoretically demonstrate their ability of robustly rejecting external disturbances (that is achieving RPA) over an exquisitely broad dynamic range. Due to their small size, flexibility, modularity, lack of side effects and applicability across various forms of life, inteins serve as promising genetic parts to implement RPA-achieving controllers. To this end, we believe “inteins in the control loop” will leave a significant impact on various disciplines spanning synthetic biology, biofuel production, metabolic engineering and cell therapy among others.

---

one of the essential features of living systems is their ability to maintain a robust behavior despite disturbances coming from their external uncertain and noisy environments. This feature, in pure biological terms, is referred to as homeostasis, which is typically achieved via endogenous feedback regulatory mechanisms shaped by billions of years of evolution. Pathological diseases are often linked to loss of homeostasis (1, 2). The need to restore homeostasis for preventing such diseases, when endogenous regulatory mechanisms fail, was one of the main drivers of ushering a new active field of research referred to as cybergenetics (3) — a field that brings control theory and synthetic biology together. In particular, the rational design and implementation of biomolecular feedback controllers (4–13) offer promising candidates that may accompany or even replace such failed mechanisms (14–16).

A notion, which is similar to homeostasis, but more stringent, is Robust Perfect Adaptation (RPA) (see e.g. (17, 18)) which is the biological analogue of the well-known notion of robust steady-state tracking in control theory. A controller succeeds in achieving RPA if it drives the steady state of a variable of interest to a prescribed level despite varying initial conditions, uncertainties and/or constant disturbances. Motivated by the internal model principle (19), which establishes that the designed controller must implement an integral feedback component to be able to achieve RPA, the antithetic integral feedback controller (20) was brought forward. The basic antithetic integral feedback motif is depicted in Fig. 1(a). It is comprised of two species **Z**_**1**_ and **Z**_**2**_ whose end goal is to robustly steer the concentration of the output species of interest **X**_**L**_ to a prescribed level, referred to as the setpoint, in spite of disturbances and uncertainties in the regulated network — represented here as the various reactions occurring between species **X**_**1**_ through **X**_**L**_. RPA is achieved via four controller reaction channels. First, **Z**_**1**_ is constitutively produced at a rate *µ* to encode for the setpoint. Second, **Z**_**2**_ is catalytically produced from the output species **X**_**L**_ at a rate *θx*_*L*_ to sense its concentration. The third reaction is the annihilation or sequestration reaction between **Z**_**1**_ and **Z**_**2**_ occurring at a rate *ηz*_1_*z*_2_. The sequestration reaction encodes a comparison operation and produces an inactive complex that has no function and thus its concentration need not be mathematically tracked. Finally, the feedback control action (actuation) is encrypted as a production reaction of the species **X**_**1**_, which acts as the input of the regulated network, at a rate *kz*_1_ proportional to the concentration of controller species **Z**_**1**_. The underlying *Ordinary Differential Equations* (ODEs) governing the dynamics of the concentrations of **Z**_**1**_ and **Z**_**2**_ are shown in Fig. 1(a). Throughout the paper, bold uppercase letters (e.g. **Z**_**1**_) denote the names of biochemical species, while their corresponding lowercase letters (e.g. *z*_1_) denote their concentrations. By looking at the dynamics of *z*_1_−*z*_2_, it is straightforward to reveal the integral control action where the temporal error *µ/θ− x*_*L*_(*t*) at time *t*, or deviation of the output concentration from the setpoint *µ/θ*, is mathematically integrated. This establishes that, as long as the closed-loop system is stable (i.e. asymptotically converges to some fixed point), the output concentration *x*_*L*_ will converge to the prescribed setpoint *µ/θ* which is independent of the regulated network and initial coditions, and thus achieves RPA. While RPA is a steady-state property, the transient dynamic properties and tuning of the antithetic integral controller are also extensively studied as well (21–23).

**Fig. 1.**
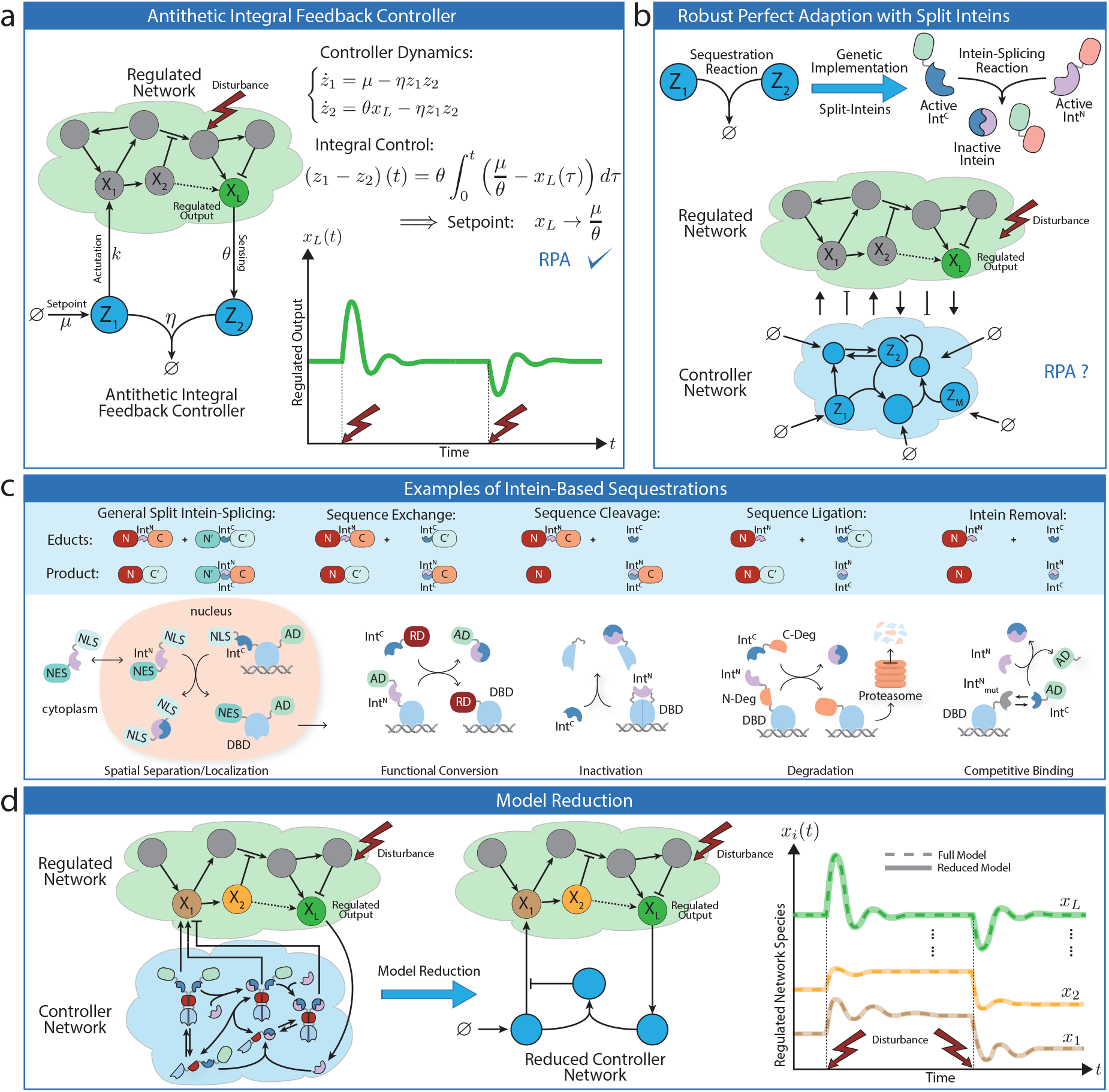
Overview. (a) The closed-loop system is comprised of the controller network (**Z**_**1**_, **Z**_**2**_) connected in a feedback configuration with an arbitrary regulated network. By examining the controller dynamics, it is straightforward to uncover the integral control action that endows the closed-loop system with RPA. That is, as long as the closed-loop system is stable, the concentration of the regulated output **X**_**L**_ converges to a prescribed value *µ/θ*, referred to as the setpoint, despite the presence of disturbances and uncertainties in the regulated network. (b) The heart of the basic AIF motif is the sequestration reaction. In this paper, we exploit the exquisite flexibility of split inteins to genetically implement a broad class of integral controllers that endow the closed-loop system with RPA. The flexibility of split inteins offers an easy-to-build biological framework at the price of (potentially) more complex mathematical models. To this end, we establish a set of simple reaction rules that enable RPA. (c) The shaded blue box schematically depicts the products of intein-splicing reactions starting from the educts. The first schematic (top left) describes the general split intein-splicing reaction where both split inteins are flanked by protein domains, labeled (N, N^*′*^) and (C, C^*′*^) for the N- and C-terminal protein sequences, respectively. The first product is a new protein containing the N and C^*′*^ domains of the educts, while the second product is a heterodimer containing N^*′*^ and C, which are held together by the two inactive split inteins. The remaining four schematics in the shaded blue box are instantiations of the general case and are labeled according to the perspective of the protein containing the Int^N^ segment. As the labels suggest, a part of the protein sequence is either exchanged for another one or removed through cleavage. Furthermore, it is possible to ligate another sequence to the protein of interest or make it non-responsive to future splicing reactions through intein removal. To illustrate the design modularity and flexibility, we list a selection of intein-based implementation examples of the antithetic sequestration motif (below the shaded blue box) based on the described possible splicing reactions. In the first example (bottom left), inteins are used to shuffle proteins between the nucleus and the cytoplasm due to the NLS and the NES which flank the protein sequences. In particular, the intein-splicing reaction exchanges the NLS with a NES which leads to the export of a TF out of the nucleus where it cannot initiate transcription anymore. In the second example, inteins are used to exchange an AD by a RD, which inverts the function of the TF. In the third example, a split inteins is introduced within a functional domain without disturbing it. The splicing reaction results in the cleavage of the domain rendering it nonfunctional. In the fourth example, a protein is fused to the first split-intein and a split-degradation tag, while the second split-intein is fused to the other half of the split-degradation tag. The splicing reaction re-ligates the degradation tag rendering it functional and capable of degrading a POI. In the final example, a DBD can reversibly heterodimerize with an AD via its split inteins. Note that the split intein on the DBD is mutated so that it cannot perform the splicing reaction upon dimerization. A separate non mutated intein is able to remove the intein from the AD through splicing. This renders the AD unable to heterodimerize with the DBD. AD: activation domain, RD: repressing domain, Int^C^: intein C, Int^N^: intein N, DBD: DNA binding domain, NLS: nuclear localization signal, NES: nuclear export signal, N-Deg: N terminus of split degradation domain, C-Deg: C terminus of split degradation domain. (d) A simple recipe is developed to reduce the otherwise mathematically complex controller models into simple motifs that resemble the basic AIF motif, but with a fundamental difference: the sequestration product is allowed to have a function that can be leveraged as a tuning knob to enhance the controller performance while maintaining RPA.

*In vivo* antithetic integral feedback controllers have been previously built in both bacteria (4, 6) and mammalian cells (7), where RPA is experimentally demonstrated. A quasi-integral controller using a slight variant of the antithetic controller was also demonstrated in *E. coli* (9). More recently, a protein based antithetic controller in mammalian cells was also recently proposed (24). In (6), the controller is implemented in *E. coli* using sigma/anti-sigma factors as the basic parts that realize the sequestration reaction — the heart of the antithetic integral motif. In (7), the controller is implemented in HEK293T cells using sense/anti-sense mRNAs. The sequestration reactions in both designs are achieved by the heterodimerization of **Z**_**1**_ and **Z**_**2**_. In the case of sigma/anti-sigma factors, the heterodimerization reaction is reversible, a fact that may lead to reduced performance in certain operating regimes. Moreover, when present in high quantities, xenogenic sigma factors may be toxic due to their inherent property of sequestering RNA polymerases from housekeeping sigma factors (25). Finally, sigma factors are specific to the transcription mechanism of bacteria and cannot be easily transferred to other domains of life, which is why the sense/anti-sense RNA controller (7) was developed for mammalian cells. At the same time, sense/anti-sense hybridizations produce double-stranded RNAs that, in high abundance, may initiate global translational repression (26), leading to a reduction in the effective dynamic range of operation. These constraints give rise to the need for genetic parts that are nontoxic, are transferable between different forms of life, and enjoy wider dynamic ranges of operation. Nevertheless, the suitable choice of genetic parts is a difficult task because they need to adhere to the strict design rules of the basic antithetic integral feedback motif. In this paper, we show that split inteins serve as the ideal candidate parts that are capable of doing both: adhering to the design rules and avoiding the aforementioned disadvantages. We build on the universality result of the antithetic motif (6) to examine more complex integral controller designs for RPA as demonstrated in Fig. 1(b). While more complex mathematical topologies do not have to be necessarily more difficult to implement, they certainly broaden the biological design space. This expansion, in fact, becomes necessary due to the biological implementation constraints.

An intein is a protein segment that is capable of autocatalytically excising itself from the protein while religating the remaining segments, called exteins, via forming new peptide bonds (27). Inteins are universal as they can be naturally found in all domains of life spanning eukaryotes, bacteria, archaea and viruses (28, 29). Split inteins — a subset class of inteins — are, as the name suggests, inteins split into two halves commonly referred to as Int^N^ and Int^C^. Split inteins have been widely studied and characterized due to their extensive usage in various life science disciplines and their ability to perform fast, reliable and irreversible post-translational modifications (30–34). Small split inteins like Gp41-1^C^ are comprised of around 40 amino acids (35) and are well within the size range of synthetic protein linkers (36). It is then possible to use them as “functional” linkers to connect different protein segments. The split inteins, when active, are capable of heterodimerizing and performing protein splicing reactions on their own where they irreversibly break and form new peptide bonds in a strict stoichiometric ratio of one to one. We shall refer to these reactions as “intein-splicing reactions” where molecules containing active Int^C^ segments react with molecules containing active Int^N^ segments to undergo a particular splicing mechanism. When two molecules undergo an intein-splicing reaction, the Int^N^ and Int^C^ segments are permanently inactivated as they are unable to perform further splicing reactions due to the alteration of their respective biochemical structures. However the products of such a reaction may still have other functions such as activating or repressing gene expression due to the presence of other protein domains that may not be affected by the splicing reaction. Split inteins can be exploited to exchange, cleave or ligate amino acid sequences (see Fig. 1(c)). These features serve as the basis of realizing the sequestration reaction of the antithetic integral motif. A selection of antithetic “sequestrations” based on functional conversion, spatial separation, inactivation, degradation and intein removal are shown in Fig. 1(c) to emphasize the modularity and the vast flexibility of intein-based designs. Nonetheless, this high design flexibility comes with a price: simple intein-based implementations may lead to complicated network topologies very quickly as illustrated in Fig. 1(d). Here we exploit a time-scale separation argument to establish a structural model reduction result which provides an easy-to-use recipe to simplify the underlying models. This facilitates the mathematical analysis of the otherwise complicated controller network, and allows us to uncover the underlying controller structure which is not necessarily limited to integral control only.

Integral control is the fundamental building block in most controllers spanning a broad range of industrial applications in the fields of electrical, mechanical and chemical engineering; however, it is rarely used alone. In fact, Integral (I) controllers are typically augmented with Proportional (P) and/or Derivative (D) controllers to obtain PI/PID controllers that offer more flexibility in enhancing the dynamic performance while maintaining the RPA property. Recently, more advanced molecular controllers such as PI/PID controllers found their way to molecular biology (7, 37–42). Ideally, pure proportional control is achieved via instantaneous negative feedback from the output **X**_**L**_ to the input species **X**_**1**_ and it is shown that it is not only capable of enhancing the transient dynamic performance, but also reducing cell-to-cell variability (37, 40). The first biomolecular (filtered) PI controller was genetically engineered in (7) where additional genetic parts are appended to the antithetic integral motif to realize the proportional component. Here, we establish that a filtered PI controller can be built without introducing additional genetic parts by harnessing the sequestration products of the split inteins (38).

Besides proposing new intein-based implementation strategies for RPA-achieving controllers and laying down the necessary theoretical foundation, we have also selected, built and tested five structurally different controller topologies for experimental verification of RPA. All circuits were tested in HEK293T cells and range from pure I to filtered PI controllers based on the functional conversion, inactivation and intein removal strategies illustrated in Fig. 1(c).

## Results

Split inteins offer a high degree of flexibility in realizing biomolecular integral feedback controllers. This flexibility is mainly a consequence of their compatibility with essentially any transcription factor. In fact, the particular structure of the expressed transcription factor including the choices of the Activation Domain (AD), Dimerization Domain (DD), DNA-Binding Domain (DBD) and insertion position of the split intein (Int^C^) open the possibilities to a broad design space of controllers. Specifically, dimeric transcription factors, such as tTA, give rise to multiple homo- and hetero-dimerization reactions as well as multiple sequestration reactions and thus make the controller network more complex to mathematically analyze. To this end, we develop a theoretical framework tailored to mathematically analyze and simplify complex intein-based controller networks that generalize the basic antithetic integral motif which has no dimerization reactions and a single sequestration reaction.

## Achieving Robust Perfect Adaptation using Inteins

In this section, we establish a theoretical framework embodied as a set of simple rules that allows us to design biomolecular controllers enabling RPA using split inteins. Consider the general closed-loop network depicted in Fig. 2 where an arbitrary network comprised of *L* species **X** := {**X**_**1**_, …, **X**_**L**_}, referred to as the regulated network, is in a feedback interconnection with the controller network comprised of *M* species **Z** := {**Z**_**1**_, …, **Z**_**M**_}. The overall objective of the feedback controller network is to achieve RPA of the regulated output species **X**_**L**_ by *automatically* actuating (producing and/or degrading) the input species **X**_**1**_. Each controller species **Z**_***i***_, for *i* = 1, 2, …, *M*, belongs to one of three classes: 𝒞 -class, 𝒩 -class and 𝒮-class. These classes separate the controller network into three subnetworks as depicted in Fig. 2. The classification of the controller species and the allowed reactions follow the rules that are listed in Fig. 2. In particular, the setpoint and sensing of the regulated output species **X**_**L**_ are encoded in the constitutive and/or catalytic production reactions following Reaction Rule 1 given by

**Fig. 2.**
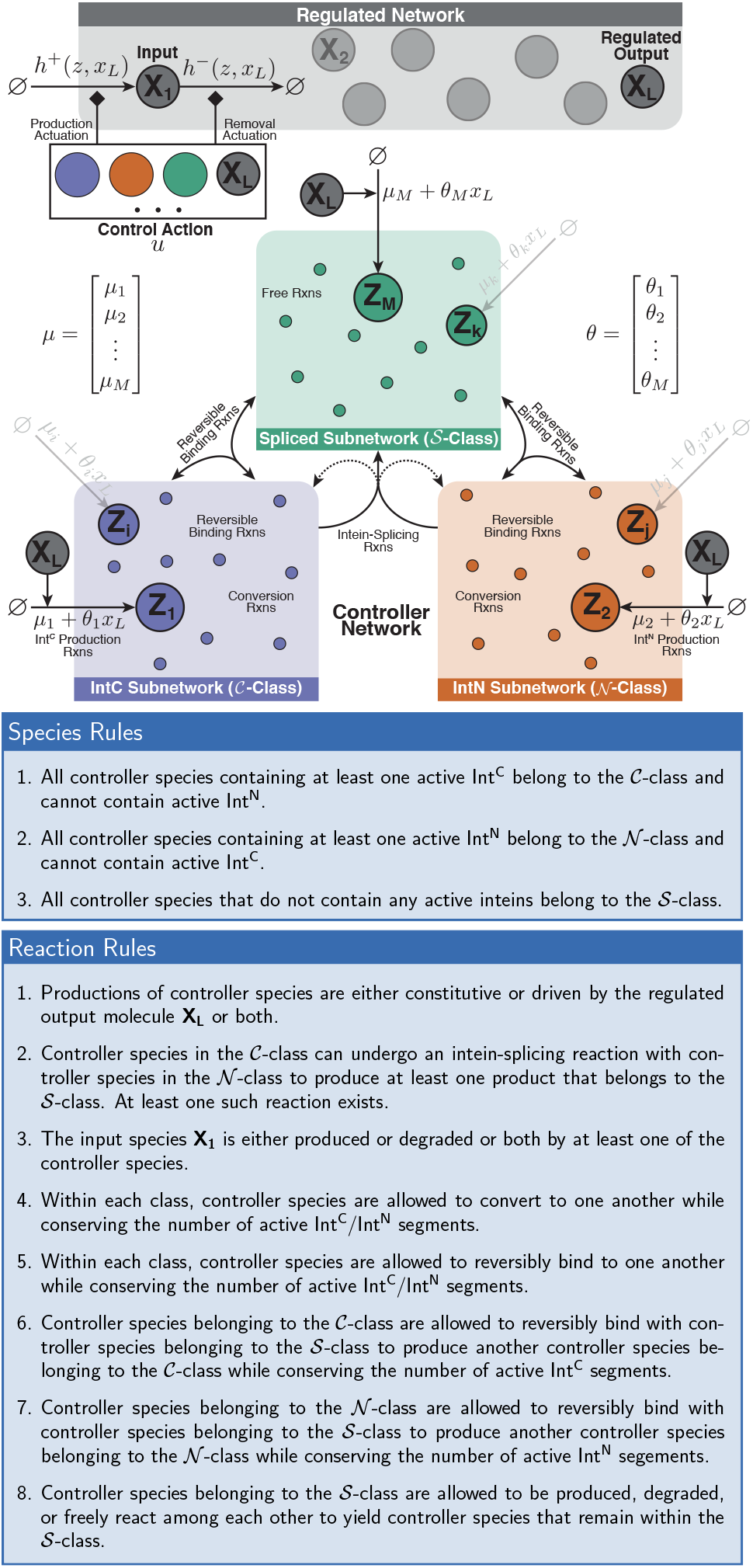
A theoretical framework for RPA-achieving intein-based integral controllers. The closed-loop network is formed of a controller network, comprised of *M* species **Z**_**1**_, …, **Z**_**M**_, connected in a feedback configuration with the regulated network, comprised of *L* species **X**_**1**_, …, **X**_**L**_. Following the general biomolecular control paradigm (37), it is assumed that the controller interacts with the regulated network via **X**_**1**_ and **X**_**L**_ only, referred to as the input and regulated output species, respectively. The objective of the controller network is to steer the concentration of the regulated output **X**_**L**_ to a prescribed value, referred to as the setpoint, despite the presence of constant disturbances and uncertainties in the regulated network, that is, ensuring RPA of **X**_**L**_. The controller network is divided into three subnetwork classes according to the list of Species Rules, and the allowed reactions within and between the three subnetworks are listed as Reaction Rules. The feedback controller network operates by “sensing” the abundance of the concentration of the regulated output **X**_**L**_ (*θ*_*i*_x_*L*_), and “actuating” the input **X**_**1**_ by producing it (*h*^+^(*z, x*_*L*_)) or removing it (*h*^−^(*z, x*_*L*_)). The total control action *u* is given by *h*^+^(*z, x*_*L*_) *− h*^−^(*z, x*_*L*_)*x*_1_. Note that, throughout the paper, the diamond-shaped arrowhead denotes either an activation or repression. The setpoint and output-sensing mechanisms are jointly encoded in the vectors *µ* and *θ* to allow for multiple setpoint-encoding reactions.

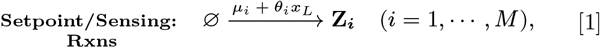

with at least one *µ*_*i*_ and one *θ*_*i*_ strictly positive. The following theorem provides a guarantee for RPA of the regulated output species when controlled with intein-based controllers.

### Theorem 1 (RPA for Intein-Based Controllers)

*Consider the closed-loop network depicted in Fig. 2 where the controller network respects the set of listed rules. Let* 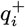 *and* 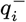 *respectively denote the number of active Int*^*C*^ *and Int*^*N*^ *segments present in controller species* **Z**_***i***_ *for i* = 1, …, *M*. *Define* 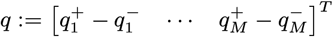. *Then, if the closed-loop network is stable, the controller network ensures RPA of* **X**_**L**_ *with*

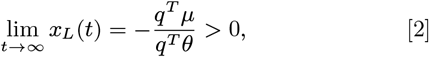

*where µ* :=[*µ*_1_ *… µ*_*M*_]^*T*^ *and θ* := [*θ*_1_ *… θ*_*M*_]^*T*^. *Furthermore, the integral variable is given by z*_*I*_ := *q*^*T*^ *z which reveals the underlying integral controller given by*

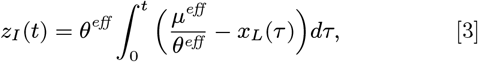

*with µ*^*eff*^ := *q*^*T*^ *µ and θ*^*eff*^ := *−q*^*T*^ *θ*.

Before we proceed, we provide two remarks.

### Remark 1.1

Theorem 1 is a special case of a more general theorem which can be also applied to any non-intein-based biomolecular controller with similar structure as demonstrated in the example of Box 1. This more general theorem interprets *q*^+^ and *q*^−^ as the number of positive and negative charges (where, here, the number of inteins is an instantiating of the charge analogy) and extends the RPA sufficiency result in (6) to the case of multiple sensing and setpoint reactions. In fact, if **Z**_**1**_ is the only controller species that is constitutively produced and **Z**_**2**_ is the only controller species that is catalytically produced by the regulated output species **X**_**L**_, then *µ*_1_, *θ*_2_ *>* 0, *µ*_*i*_ = *θ*_*j*_ = 0 for (*i, j*)*≠* = (1, 2) and *q* = [1 −1 *⋆*] ^*T*^ which yields 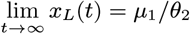 — the RPA result in (6).

### Remark 1.2

The catalytic sensing terms *θ*_*i*_*x*_*L*_ for *i* = 1, …, *M* shown in Fig. 2 and Eq. (1) do not necessarily have to be linear. In fact, these terms can be replaced by more general nonlinear functions *f*_*i*_(*x*_*L*_) such as Hill-type functions that are allowed to be monotonically decreasing to account for repressive sensing. These sensing mechanisms will preserve RPA in the deterministic setting, but the setpoint expression will be different from Eq. (2).

### Implementations using Various Transcription Factors

So far, we have described, theoretically, how split inteins can be exploited to build a broad class of biomolecular integral controllers capable of achieving RPA. Here, we demonstrate how commonly used transcription factors can be converted into controller species that respect the rules of Fig. 2 and, as a result, enables RPA according to Theorem 1. In particular, we use three common DNA binding domains: ZF (43, 44), TetR (45) and Gal4 to construct four structurally different biomolecular controllers (Fig. 3), that serve as instantiations of the class of controllers described in Theorem 1. We also provide experimental proof (Fig. 4) that these intein-based controllers are indeed capable of achieving RPA and thus rejecting perturbations over a wide dynamic range.

**Fig. 3.**
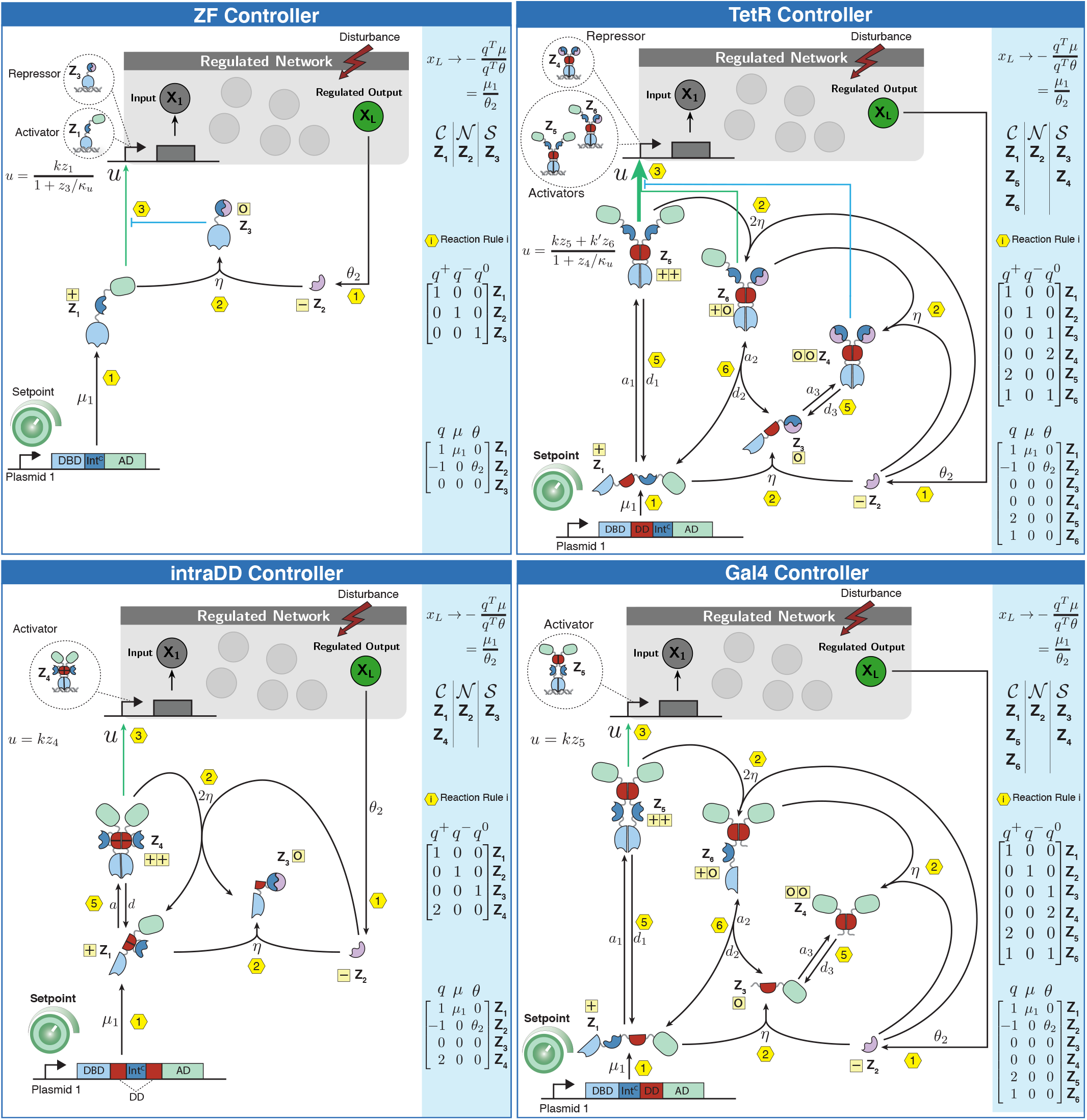
Intein-based implementation of RPA-achieving integral controllers using ZF, TetR and Gal4 as DBDs. In all four circuits, the protein **Z**_**1**_ is constitutively expressed at a rate *µ*_1_ from Plasmid 1 to encode for the desired setpoint. One of the two main tasks of **Z**_**1**_, which contains one Int^C^ within a TF, is to either directly actuate the regulated network by producing the input species **X**_**1**_, or to dimerize first and then perform the actuation. The second main task of **Z**_**1**_ is to undergo an intein-splicing reaction with the second split intein Int^N^, denoted by **Z**_**2**_, whose production is driven by the regulated output **X**_**L**_ at a rate *θ*_2_ *x*_*L*_ to encode for the “sensing” reaction. Different positions of the Int^C^ segment and different TF structures give rise to different control topologies. When controller species contain a DD, reversible homo- or hetero-dimerization reactions occur with an association rate *a*_*i*_ and a dissociation rate *d*_*i*_; whereas, when a controller species containing an active Int^C^ segment meets with another controller species containing an active Int^N^ segment, an irreversible intein-splicing reaction occurs at a rate *η* multiplied by an integer that depends on the number of participating inteins. Note that the splicing products which do not have any activity are omitted here for simplicity. All controller species directly impacting the regulated network are indicated either as repressors or activators in the dashed bubbles, and the mathematical expression of the total control action *u* is shown as a (Hill-type) function of the repressors and activators. Every reaction is labeled from 1 to 6 according to the permitted reaction rules stated in Fig. 2. Furthermore, every monomer, independent of its dimerization status is labeled in the yellow boxes with one of the following “charges”: +,−, 0, according to Theorems 1 and 2. This is also repeated in the charge vectors *q*^+^, *q*^−^ and *q*^0^ that encode, for each controller species, the number of active Int^C^, Int^N^, and monomers with no active inteins (see Theorem 1 and 2). Furthermore, in the blue boxes, all controller species are grouped into *C, N* and *S* classes according to the split inteins they contain (see Species Rules in Fig. 2). Since all the Species and Reaction Rules of Fig. 2 are respected, then by Theorem 1, we conclude that all four controllers ensure RPA (as long as closed-loop stability is maintained), and the setpoint can be shown, using Eq. (2), to be *µ*_1_ */θ*_2_.

**Fig. 4.**
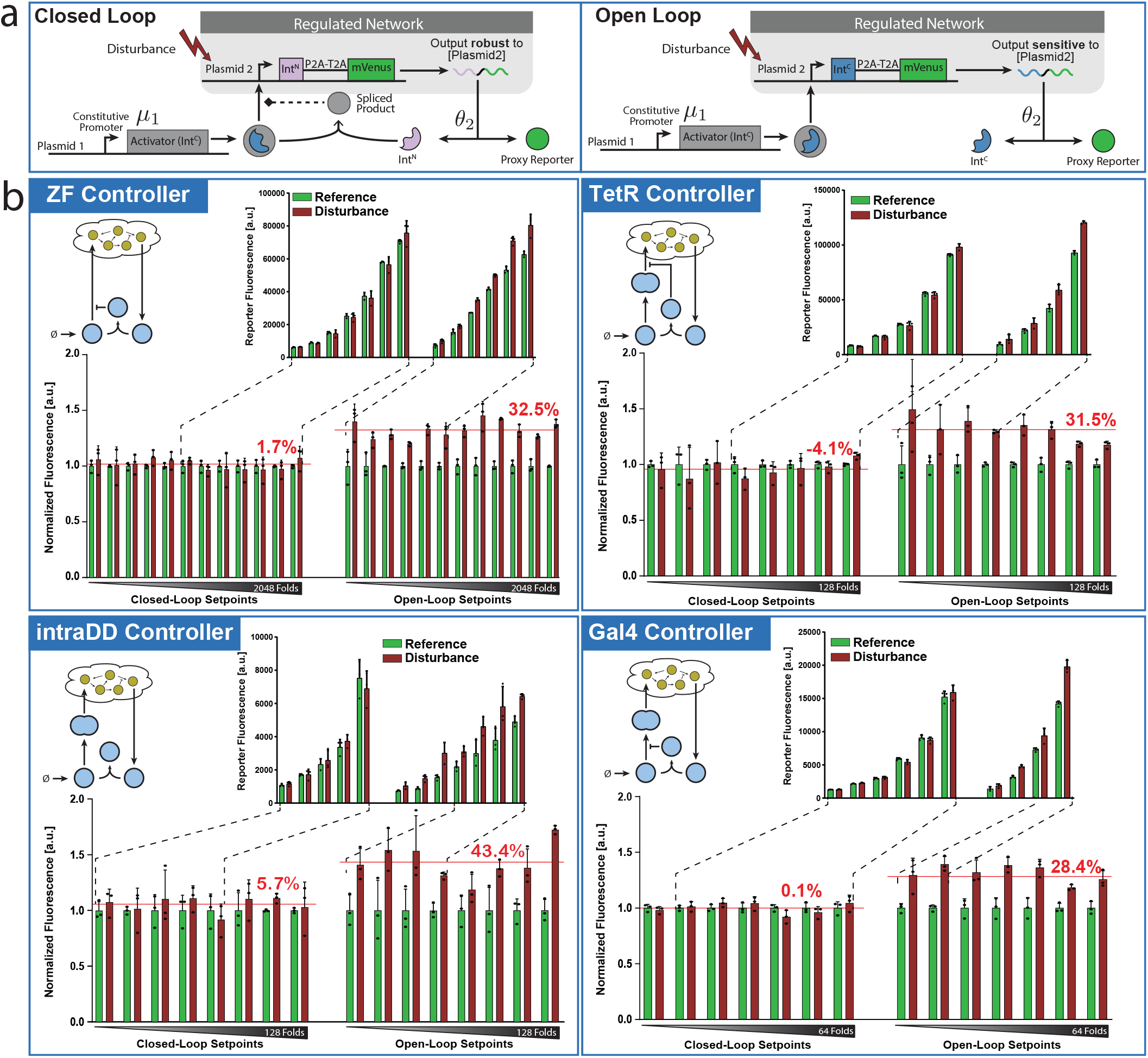
Experimental Demonstration of RPA. (a) Illustration of the experimental setup for closed-loop characterization. All four controllers of Fig. 3 were tested in a two-plasmid system. Plasmid 1 encodes an Int^C^ segment incorporated in an activator which is driven by a constitutive promoter. Plasmid 2 encodes either Int^N^ (in the closed-loop setting) or Int^C^ (in the open-loop setting), which is fused to the fluorophore mVenus via a P2A-T2A linker and is driven by the activator expressed from Plasmid 1. Note that the P2A-T2A linker leads to the translation of two separate proteins (Int^N^ and mVenus) in a fixed ratio from a common mRNA due to ribosome skipping (51). The fluorescent protein mVenus is used as a proxy reporter for its own mRNA which is the regulated output that is expected to exhibit RPA when operating in closed loop. In the open-loop setting, there is no interaction between the two Int^C^ segments as they are unable to homodimerize and cannot perform the intein-splicing reaction. However, in the closed-loop setting, the pair can perform the intein-splicing reaction to produce further products which are, in certain cases, capable of directly acting on the regulated network such as repressing the transcription of the gene in Plasmid 2. The setpoint is tuned by changing the constitutive production rate *µ*_1_ via the transfected copy numbers of Plasmid 1. The controller “senses” the abundance of the regulated output via the translation of the mRNA encoded in Plasmid 2, whose rate is denoted by *θ*_2_. Finally, Plasmid 2 is used to apply a disturbance to the regulated network because its concentration does not alter the setpoint given by *µ*_1_ */θ*_2_. In fact, the reference, or undisturbed output level, is obtained by fixing the copy number of the transfected Plasmid 2 to a certain value across all setpoints; whereas, the disturbed output level is obtained by repeating the same experiment for all setpoints but with a higher copy number of Plasmid 2. (b) Steady-state errors. For each controller circuit, a simplified schematic (top left) of the controller topology is shown, and two bar graphs are sketched to report the experimental measurements. The bottom bar graphs show the normalized fluorescence of the proxy reporter with (disturbance, red) and without (reference, green) disturbance and for both the closed-loop (left) and open-loop (right) settings. The disturbed and reference triplicate measurements were normalized to the mean fluorescence of the reference data for each setpoint. The x-axis follows a log_2_ -scale and shows the amount of Plasmid 1 transfected within every well. The red horizontal lines give the normalized output averaged over all the setpoints, and the numbers above the lines indicate the averaged error of the disturbed output relative to the reference. The top figures, for each controller circuit, show the non-normalized data over a selected subset of setpoints that match in absolute fluorescence between the open- and the closed-loop circuits. The dashed lines point to the selected setpoints which were used for matching the output fluorescence.

There are a few considerations that have to be taken into account to successfully build intein-based integral controllers.

Building Consideraion

1. Fusion of the split intein to or into a protein of interest should not significantly impair the function of the target.
2. The split intein has to be able to fold properly to efficiently perform the intein-splicing reaction.
3. The split intein has to be sterically accessible by its other split intein to be capable of efficiently performing the intein-splicing reaction.

These considerations should be experimentally verified for any intein-based controller to function properly. In particular, to minimize the impairment of the protein of interest as per Building Consideration 1, we use the smaller Int^C^ of the fast reacting intein Gp41-1 (46) for all our modified activators.

Next, we provide a detailed description of the four different controller circuits depicted in Fig. 3. We start with the ZF controller which has the simplest topological structure. It is obtained by using ZF as the DBD, and introducing the split intein in the floppy linker between the AD VP64 and the DBD ZF. This TF, denoted by **Z**_**1**_, is constitutively produced at a rate *µ*_1_ and is capable of actuating the regulated network of interest by activating the expression of the input **X**_**1**_. The regulated output **X**_**L**_ produces the second split intein Int^N^, denoted by **Z**_**2**_, at a rate *θx*_*L*_ that is proportional to the regulated output concentration. The intein-splicing reaction between **Z**_**1**_ and **Z**_**2**_ occurs at a rate *η* and leads to a cleavage within the TF, which separates the AD from the DBD. The resulting free floating AD is not tracked due to its inability to initiate transcription on its own. The other spliced product is the DBD, denoted by **Z**_**3**_, which competes with **Z**_**1**_ for the promoter binding sites, and thus exerts a repressive actuation.

The second controller design, labeled as intraDD, is based on TetR whose goal is to illustrate that it is possible to build intein-based antithetic integral controllers without functional spliced products. This controller is obtained by introducing the split intein within the DD of TetR without disrupting it. The transcription factor **Z**_**1**_ is generated by fusing VPR to the modified TetR. The dimer **Z**_**4**_ comprised of two molecules of **Z**_**1**_ acts as the actuating controller species. Unlike the previous controller, Int^N^, denoted by **Z**_**2**_, can now undergo an intein-splicing reaction with either the monomer **Z**_**1**_ or the dimer **Z**_**4**_. The intein-splicing reaction with **Z**_**1**_ leads to the cleavage of the protein sequence next to the Int^C^, which is acting as a linker holding the two halves of the split DD together. This results in two products: the AD VPR with part of the disabled DD, denoted by **Z**_**3**_, and a monomeric TetR with the rest of the disabled DD (not tracked in Fig. 3 due to its inactivity). Neither of them are able to further interact with the controller or the regulated network. Similarly, the intein-splicing reaction with the dimer **Z**_**4**_ leads to the cleavage of one of the monomers within the DD. This results in the immediate falling apart of the dimer into one **Z**_**1**_ and one **Z**_**3**_.

The third controller design is obtained by inserting an Int^C^ segment between TetR and the AD. The expressed TF, denoted by **Z**_**1**_, has to dimerize to form **Z**_**5**_ to be able to actuate the regulated network. Int^N^, denoted by **Z**_**2**_, can undergo an intein-splicing reaction with either the monomer **Z**_**1**_ or the dimer **Z**_**5**_. The intein-splicing reaction between **Z**_**1**_ and **Z**_**2**_ leads to the separation of the AD from the remaining DBD and DD to produce **Z**_**3**_ and a free floating AD which is not tracked anymore due to its inactivity. The spliced product **Z**_**3**_ can still heterodimerize with **Z**_**1**_ to yield a TetR dimer with only one AD, denoted by **Z**_**6**_, which is sufficient to bind to the promoter and initiate transcription. Note that **Z**_**6**_ can be also obtained via the intein-splicing reaction between the fully intact dimer **Z**_**5**_ and **Z**_**2**_. Furthermore, since **Z**_**6**_ still has one functional Int^C^ segment, it is able to perform a second intein-splicing reaction with **Z**_**2**_, which removes the last AD by cleavage and hence forms a tetR dimer, denoted by **Z**_**4**_. Note that, this dimer can be also obtained via the homodimerization of **Z**_**3**_. The dimer **Z**_**4**_ can recognize and bind to the promoter, but can not initiate transcription unlike the other two dimers **Z**_**5**_ and **Z**_**6**_. It therefore, competes with **Z**_**5**_ and **Z**_**6**_ for the promoter binding sites and, as a result, acts as a repressor.

The last controller design of Fig. 3 is based on the yeast derived DBD Gal4 and is thus labeled as the Gal4 controller. Here, we introduced an Int^C^ segement between the DBD and the DD. Similar to TetR, Gal4 needs to be a dimer (**Z**_**5**_) in order to bind to the promoter and actuate the regulated network. Once again, Int^N^, denote by **Z**_**2**_, can undergo an intein-splicing reaction with either **Z**_**5**_ or **Z**_**1**_. The intein-splicing reaction with **Z**_**1**_ leads to the separation of the DBD from the remaining DD and the AD to produce **Z**_**3**_. As already mentioned, Gal4 cannot bind to the promoter as a monomer, and so we do not track this species due to its inactivity. Furthermore, the intein-splicing reaction with **Z**_**5**_ leads to the removal of one DBD from the dimer through cleavage, which renders the entire complex unable of binding to the DNA. This truncated dimer, denoted by **Z**_**6**_, can perform a second intein splicing reaction with **Z**_**2**_ to remove the second DBD and form a new dimer denoted by **Z**_**4**_ which is also incapable of acting directly on the regulated network. However, it is able to disassociate into its monomers, **Z**_**3**_, which are able to reversibly sequester **Z**_**1**_ through a heterodimerization reaction yielding the nonfunctional dimer **Z**_**6**_.

It is fairly straight forward to verify that all the reaction rules listed in Fig. 2 are respected by all of the proposed four controllers. As a result, by applying Theorem 1, we conclude that all four proposed controllers achieve RPA (as long as the closed-loop network is stable) such that the concentration of the regulated output *x*_*L*_ converges to *µ*_1_*/θ*_2_ at steady state. Next, we provide an experimental verification to back up our developed theory. To do so, all of the four proposed controller circuits were first tested for the three Building Considerations. In fact, to test for Building Consideration 1, we expressed all of the modified activators constitutively and compared their ability to transcribe a fluorophore. We observed a drop in activity for all modified ZF, tetR and GAL4 based TFs ranging from significant to minor. To this end, strong impairments were partially compensated by using stronger activation domains like VPR. Intein insertions within floppy linkers were relatively straight forward; however, insertions within functional protein domains, as was the case for the intraDD-Circuit (see Fig. 3), required some screening. Next we tested for Building Considerations 2 and 3 by constitutively expressing the modified activator carrying Int^C^ together with the second split intein (Int^N^) and observed the levels of a fluorescent reporter. If the Building Considerations are satisfied the fluorescent output will decrease with increased levels of Int^N^. We were able to reach background levels for every controller type upon a high expression of the second split intein. This indicates that the intein-splicing reaction is indeed happening as expected.

After making sure that all Building Considerations were fulfilled, we proceeded with characterizing the controllers in the closed-loop setting. We opted for a simple two-plasmid, closed-loop system for testing the controller performance as demonstrated in Fig. 4(a). This allowed us to focus on the controller behavior without having to worry about potential cross-talks (47), resource burden (48) or saturation (49) which might appear in larger circuits. The first plasmid encodes for the modified transcription factor **Z**_**1**_ and the other one encodes for either Int^C^ for the open-loop circuit or Int^N^ for the closed-loop circuit. In both cases, the split intein was encoded with a P2A-T2A linker and the fluorophore mVenus. The fluorophore is used as a proxy for its own mRNA, which is the regulated species expected to exhibit RPA. The advantage of this setup is that changing the copy numbers of the two transfected plasmids can be conveniently used to characterize the controllers. More precisely, *µ*_1_ and hence the setpoint can be easily tuned by altering the amount of the plasmid encoding for the activator. Furthermore, the translation rate *θ*_2_ of the mRNA is independent from the plasmid copy numbers in the cell. Perturbing the copy numbers of plasmid 2 only leads to an increase in the transcription rate of the output mRNA and should be rejected if the integral controller works as expected. Hence, to experimentally test the four controllers for RPA, we perturb the regulated network by increasing the copy number of plasmid 2 as it does not affect the setpoint parameters *µ*_1_ and *θ*_2_. The experimental results, depicted in Fig. 4, detail the steady-state measurements of the reporter, serving as a proxy for the regulated output (mRNA) for all four controllers. The measurements were taken for all the circuits operating in both open and closed loop, with and without disturbance. All four circuits were able to reject the disturbance over a wide titration of plasmid 1, which defines the output setpoint through tuning *µ*_1_. The best performance was observed with the ZF circuit, which succeeded in rejecting the disturbances over the entire range from the detection limit to the onset of burden.

We have used so far only the split inteins of Gp41-1, and we have successfully shown the implementation of intein-based RPA-achieving integral controllers using different TFs. Many split intein pairs with different properties have been described in literature with some of them being orthogonal to each other (50). To demonstrate that intein-based integral controllers are not limited to Gp41-1, and that it is possible to have multiple orthogonal intein-based integral controllers within the same cell, we have modified our ZF controller accordingly. In particular, we exchanged the Gp41-1, for NrdJ-1 Int^C^, one of the many orthogonal split inteins characterized by Pinto et al. (50) and closed the loop with the corresponding Int^N^ of NrdJ-1. However, instead of using Int^C^ for the open-loop circuit, we used the Int^N^ that corresponds to Gp41-1. Finally, we performed the experiment with the same plasmid ratios, which was deemed suitable for the previous Gp41-1 ZF experiment. The disturbance rejection was only visible for the compatible intein pair, and the dynamic range was similar to the experiment performed with the Gp41-1 containing ZF.

### Model Reduction

The broad class of intein-based, RPA-achieving controllers introduced in Theorem 1 gives rise to a high degree of design flexibility and thus allows topologies that may possibly involve a large number of controller species **Z**_***i***_. Furthermore, these species are allowed to react among each other via multiple binding, conversion and intein-splicing reactions according to the Reaction Rules listed in Fig. 2. This possible large number of control species and reactions may lead to complex mathematical models of high dimensions whose dynamics are not easy to understand. In this section, we consider a subset of the general RPA-achieving controllers of Theorem 1 to provide a model reduction result that makes the otherwise complex dynamics more transparent and easy to analyze. Our model reduction result is structural in the sense that its validity is independent of the particular values of the rate parameters.

Consider the Species and Reaction Rules of Fig. 2 and replace Reaction Rule 8 with five additional rules given by

**Additional Reaction Rules**

9. All controller species in the 𝒞-class undergo an intein-splicing reaction with all controller species in the 𝒩-class.
10. Binding and conversion reactions are reversible and conserve the number of inactive (bound) Int^C^-Int^N^ complexes.
11. All controller species belonging to the 𝒮-class degrade at a rate *δ*_0_ *≥* 0.
12. All controller species dilute at a rate *δ ≥* 0.
13. The intein-splicing rate between **Z**_***i***_ and **Z**_***j***_ is *η*.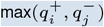 with *η >* 0.

Additional Reaction Rules

Note that Rule 9 makes Rule 2 stricter in the sense that the intein-splicing reactions are not optional anymore so that any two active intein pairs have a strictly positive propensity to undergo an intein-splicing reaction. Rule 12 takes into account the more realistic situation where *δ >* 0 which implies that RPA is not exact anymore; however, robust adaptation remains practically satisfactory as long as the dilution rate is small compared to the other rates in the network (see (6, 52)).

Finally, Rule 13 relates the intein-splicing rate to the number of participating active inteins. The following theorem provides a recipe for model reduction of (possibly complex) intein-based controllers. The model reduction result is valid in both the ideal (*δ* = 0) and non-ideal (*δ >* 0) settings and for any rate-parameter regimes.

#### Theorem 2

(Model Reduction for Intein-Based Controllers) *Consider the closed-loop network depicted in Fig. 2 where the controller network respects Species Rules 1-3 and Reaction Rules 1-7,9-13. Let* 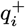 *and* 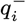 *respectively denote the number of active Int*^*C*^ *and Int*^*N*^ *segments present in controller species* **Z**_***i***_ *for i* = 1, …, *M. Let* 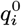 *denote the number of monomers in species* **Z**_***i***_ *with no active inteins, and construct the three vectors* 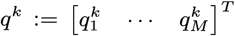, *for k ∈ {*+, −, 0*}. Furthermore, let* (*S*_*B*_, *S*_*C*_) *and* (*λ*_*B*_ (*z*), *λ*_*C*_ (*z*)) *respectively denote the stoichiometry matrices and total propensity functions associated with the reversible binding and conversion reactions that are assumed to be fast enough. If the following conditions are satisfied:*

- *S*_*B*_ *is full-column rank*.
- *The columns of S*_*C*_ *are linearly independent from those of S*_*B*_.
- *p* + rank(*S*_*C*_) = *M −* 3,

*where p is the number of reversible binding reactions, then all controller networks respecting the structure described in Fig. 2 reduce to the simple motif, depicted in Fig. 5, which is governed by only three effective species* **Z**^**+**^, **Z**^**–**^ *and* **Z**^**0**^ *whose concentrations are linear combinations of the controller species* **Z**_***i***_ *for i* = 1, …, *M*.

**Fig. 5.**
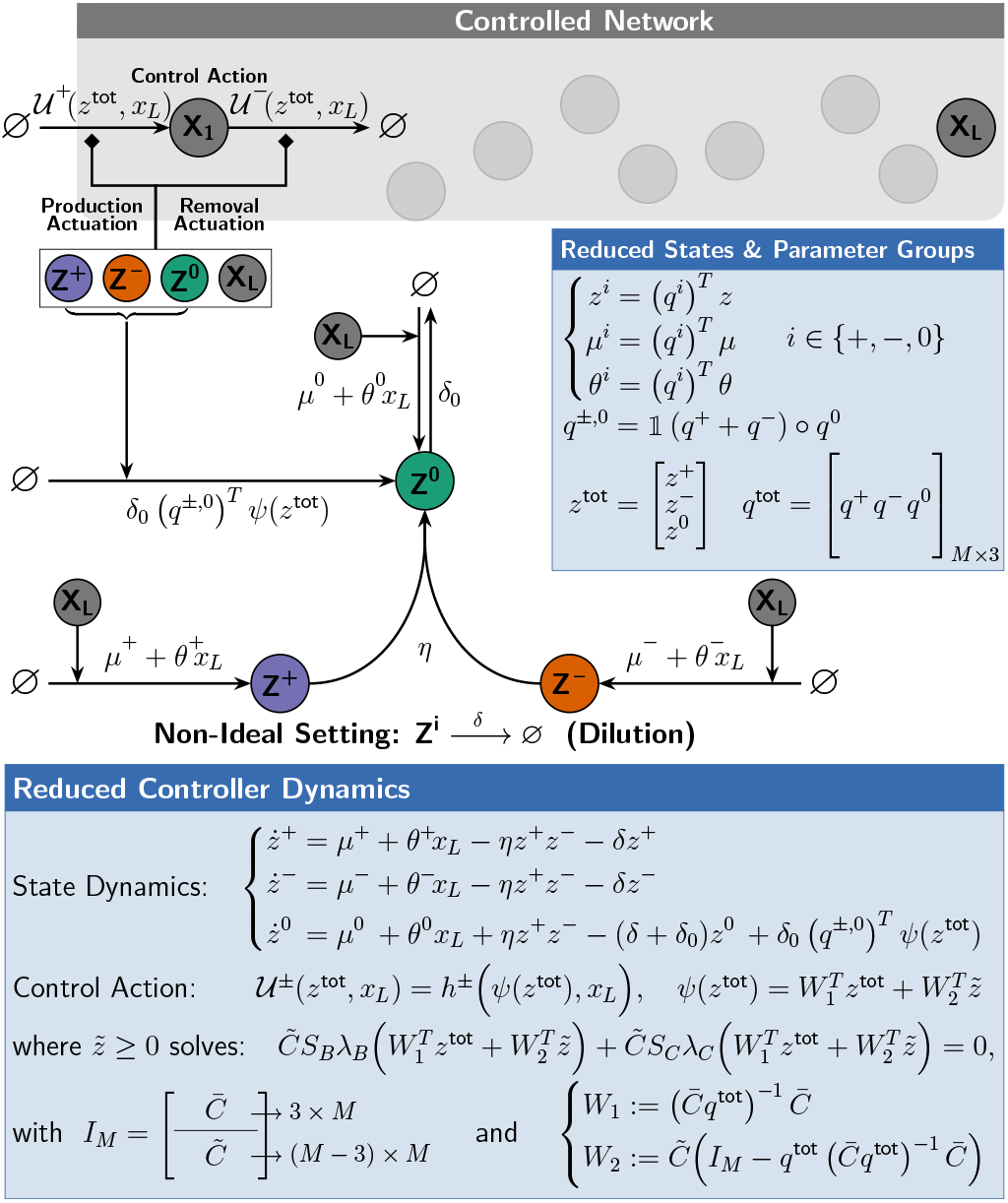
A model reduction recipe for Intein-based controllers. Under the conditions of Theorem 2, all controllers comprised of *M* species (where *M* can be large) that respect the flexible structure depicted in Fig. 2, reduce to the simple motif shown here. The reduced model is shown schematically as a motif comprised of only three effective species (**Z**^**+**^, **Z**^**–**^, **Z**^**0**^), and mathematically as a set of *Differential Algebraic Equations* (DAEs) comprised of only three differential equations in (*z*^+^, *z*^−^, *z*^0^) and *M −* 3 algebraic equations in 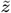. Note that (*S*_*B*_, *λ*_*B*_) and (*S*_*C*_, *λ*_*C*_) denote the stoichiometry matrices and total propensity functions (forward minus backward) of the reversible binding and conversion reactions, respectively. Furthermore 𝟙. (.), *°, I*_*M*_ and (.)^*T*^ denote the indicator function, the Hadamard (element-wise) product, the identity matrix of size *M* and the transpose of a matrix, respectively. In certain scenarios (see Fig. 6), the algebraic equations can be solved explicitly and thus further simplifying the model to only three *Ordinary Differential Equations* (ODEs). Observe that the schematic of the simple motif is fully determined once the three vectors *q*^+^, *q*^−^, *q*^0^ and the function *ψ*(*z*^tot^) are calculated. The vectors *q*^+^, *q*^−^ and *q*^0^ are easily calculated by counting active split inteins (see Theorem 2); whereas, *ψ*(*z*^tot^) can be calculated by solving the algebraic equations for 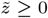 as a function of *z*^tot^.

Before we proceed, we provide four remarks.

**Remark 2.1** Once again, Theorem 2 is a special case of a more general theorem which can be also applied to any non-intein-based biomolecular controller with similar structure as demonstrated in Box 1. The proof essentially invokes the deficiency-zero theorem(53) and singular perturbation theory(54).

**Remark 2.2** The dynamics of the reduced model are depicted in the box of Fig. 5, in general, as a set of *Differential Algebraic Equations* (DAEs) comprised of only three differential equations (describing the basic effective motif) and a set of *M −*3 algebraic equations that should be solved for *z*∼*geq*0. In certain cases, these algebraic equations can be explicitly solved and thus further reducing the dynamics to a set of three ODEs (see Fig. 6). Otherwise, the algebraic equations can be left in their implicit form.

**Fig. 6.**
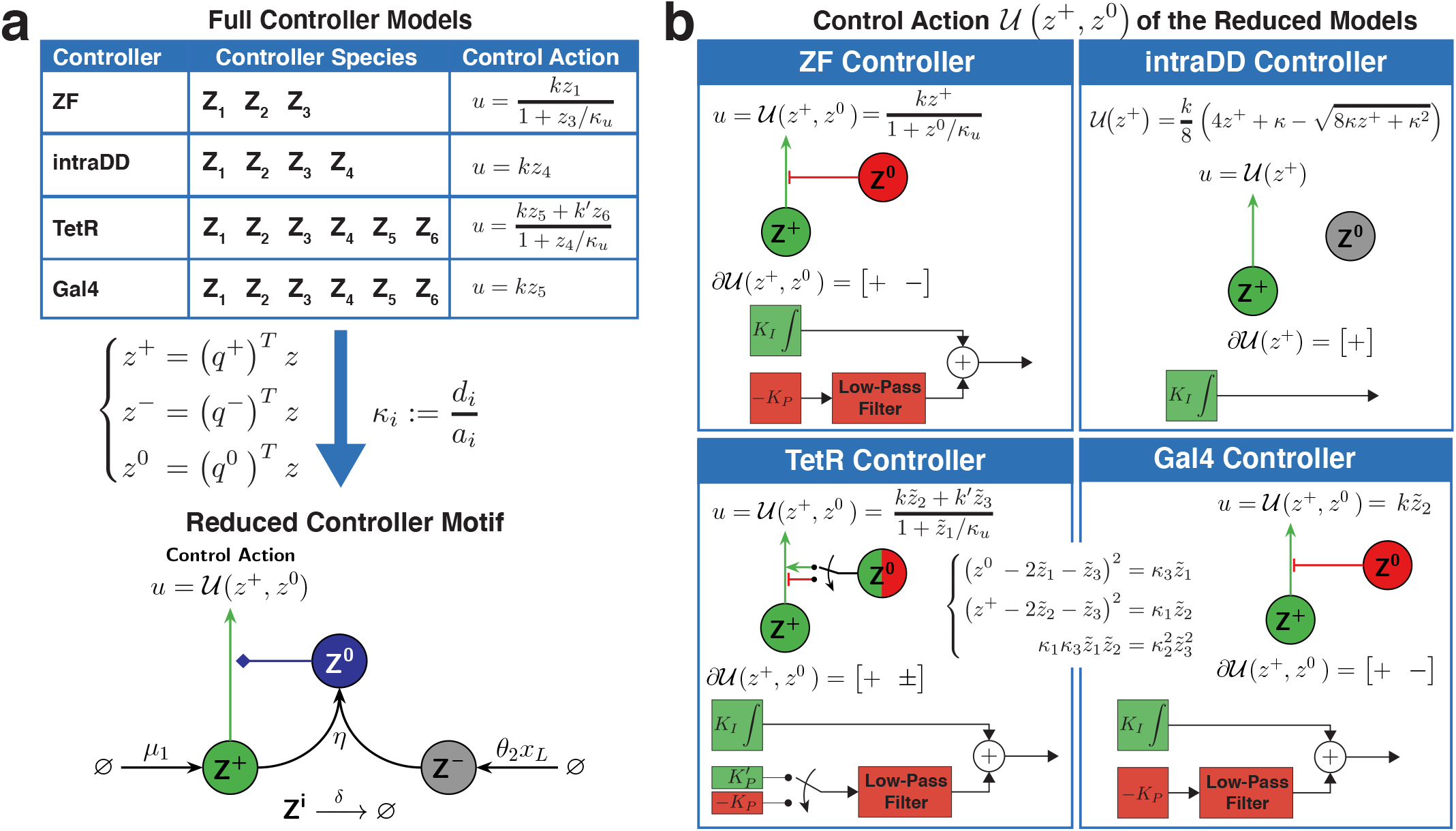
Reduced models for the ZF, intraDD, TetR and Gal4 controllers. (a) Reduced Motif. For simplicity, we assume that the protein degradation rates are negligible compared to the dilution rate *δ*; however, this assumption can be easily relaxed. Note that *δ* is assumed to be non-zero here to capture the more realistic scenario. The model reduction recipe presented in Theorem 2 and Fig. 5 can be straightforwardly applied to all of the four controller topologies in Fig. 3, where the “charge” vectors *q*^+^, *q*^−^ and *q*^0^ are shown explicitly. Observe that all four controllers reduce to the same motif comprised of the three effective species **Z**^**+**^, **Z**^**–**^ and **Z**^**0**^. The difference between them appears only in the effective control action *u* = *𝒰(z*^+^, *z*^0^). (b) Effective control actions. The control actions *u* = *𝒰(z*^+^, *z*^0^) is given separately for each controller as a function of the effective species concentrations. For the intraDD controller, the control action *u* is a strictly monotonically increasing function of *z*^+^ only, and hence the control structure is a standalone integrator. In contrast, for the ZF and Gal4 controllers, it can be shown that the control action *u* is strictly monotonically increasing (resp. decreasing) in *z*^+^ (resp. *z*^0^). This gives rise to a filtered proportional-integral (PI) control structure (38). Finally, for the TetR controller, it can be shown that the control action *u* is stricly monotonically increasing in *z*^+^; whereas, its monotonicity switches from increasing to decreasing as the levels of (*z*^+^, *z*^0^) rise. This gives rise to a filtered PI control structure where the P-component switches sign. Note that the algebraic equations presented in Fig. 5 are solved explicitly for the ZF and intraDD controllers; however, they are kept in their implicit form for the TetR and Gal4 controllers.

#### Remark 2.3.

Unlike the effective species **Z**^**+**^ and **Z**^**–**^, **Z**^**0**^ has an extra production term, in general, that is equal to *δ*_0_ [**𝟙(***q*^+^ + *q*^−^) ?*q*^0^] ^*T*^ *ψ*(*z*^tot^), where **𝟙** (.) is the indicator function, is the Hadamard (elementwise) product and *ψ*(*z*^tot^) is given implicitly in Fig. 5. This production term is zero in two cases: (1) if there are no degradation reactions (*δ*_0_ = 0), or (2) if no controller species simultaneously hold both an active intein and a monomer with no active inteins (**𝟙(** *q*^+^ + *q*^−^*)O q*^0^ = 0). Intuitively, this extra production term can be explained as follows. Controller species holding both an active intein and a monomer with no active inteins belong to either the *𝒞* - or *𝒩*-class (Species Rules), and are thus not allowed to degrade (Reaction Rules). Nevertheless, these species are still represented within **Z**^**0**^ since they hold monomers with no active inteins. As a result, the extra production term compensates for those species that do not degrade yet are represented by **Z**^**0**^ which degrades at a rate *δ*_0_.

#### Remark 2.4.

Observe that no matter what the original controller network in Fig. 1 is and as long as it satisfies the conditions of Theorem 2, the underlying effective motif is the same and is dictated by the three effective species **Z**^**+**^, **Z**^**–**^ and **Z**^**0**^ as depicted in Fig. 5. However, different controller networks give rise to different actuation functions *u*^*±*^ and production functions *ψ*. The forms of these functions lead to different control designs that may offer different tuning knobs capable of enhancing the overall performance.

Next, we apply Theorem 2 to the four controller circuits of Fig. 3 to obtain a reduced mathematical model for each. Here, we consider the more practical scenario where all controller species dilute at a rate *δ >* 0. Furthermore, we assume, for simplicity, that the degradation of the various proteins are negligible compared to the dilution rate; however, this assumption can be easily relaxed. The model reduction results are compactly depicted in Fig. 6 for all four controllers. The under-lying reduced motif, as illustrated in Fig. 6, is the same for all four controller circuits and is comprised of only three effective species **Z**^**+**^, **Z**^**–**^ and **Z**^**0**^ whose concentrations are linear combinations of the biological species **Z**_***i***_. The differences between the reduced models of each controller circuit is encrypted in the effective control action *u* = *𝒰* (*z*^+^, *z*^0^) which is a function of the concentrations of **Z**^**+**^ and **Z**^**0**^. Observe that the control action is given in an explicit form for the ZF and intraDD controllers; whereas, for the TetR and Gal4 controllers, it is given implicitly as a set of three algebraic equations. Once these algebraic equations are solved for 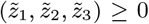 *≥* 0, the control actions can be directly computed as a function of *z*^+^ and *z*^0^. The topology of the reduced models is clearly simpler to analyze compared to the full models described in Fig. 3, and thus the underlying control architecture can be uncovered more easily. In fact, the intraDD controller realizes a standalone antithetic integral controller since the control action *u* = *𝒰* (*z*^+^) depends (monotonically) on **Z**^**+**^ only. On the other hand, it can be shown in that the control action *u* = *𝒰* (*z*^+^, *z*^0^) of the ZF- and Gal4-Circuits depends on both **Z**^**+**^ and **Z**^**0**^, such that *𝒰* is monotonically increasing (resp. decreasing) in *z*^+^ (resp. *z*^0^). This particular topology can be shown to realize a filtered Proportional-Integral (PI) controller, where the proportional component can be used as an additional knob to enhance the dynamic performance (see (38) for a thorough analysis). Finally, it can be shown that the control action *u* = *𝒰* (*z*^+^, *z*^0^) of the TetR controller also depends on both **Z**^**+**^ and **Z**^**0**^. Nevertheless,𝒰 is a monotonically increasing *𝒰* function of *z*^+^, but its monotonicity switches from increasing (at low levels of *z*^0^ and *z*^+^) to decreasing (at higher levels of *z*^0^ and *z*^+^). Interestingly, this architecture realizes a filtered PI controller whose proportional component switches from positive to negative gain. This gives rise to a nice feature that initially speeds up the response when the concentrations of the controller species are low, and then switches to negative feedback as the concentrations rise and thus favoring closedloop stability. The various reduced models are validated via simulations (not shown here) that demonstrate the highly accurate matching between the dynamics of the full and reduced models.

#### Generality of Theorem 1.

In this section, we demonstrate that Theorem 1 can be applied to controller circuits that are more general compared to those of Theorem 2. That is, there are certain intein-based controllers that can be easily tested for RPA using Theorem 1; however, their model reduction cannot be carried out by applying Theorem 2. We do so by considering the circuit depicted in Fig. 7(a). In this circuit, we constructed two genes encoding for an AD fused to an active Int^C^ (expressing **Z**_**1**_) and a DBD-DD fused to an inactive Int^N^ (expressing **Z**_**4**_). Although the inactive Int^N^ lacks essential amino acids to undergo the intein-splicing reaction (55), **Z**_**4**_ can still reversibly bind to **Z**_**1**_ to form a heterodimeric transcription factor. In this controller design, the intein-splicing reaction can occur only between the expressed Int^N^, denoted by **Z**_**2**_, and **Z**_**1**_, because **Z**_**1**_ is the only controller species that contains an active Int^C^ segment in its unbound state. In fact, although the other controller species containing active Int^C^ segments (**Z**_**6**_, **Z**_**7**_ and **Z**_**8**_) belong to the -class, they cannot directly undergo inteinsplicing reactions since they are bound to the inactive Int^N^. This results in a violation of Reaction Rule 9 rendering the model reduction recipe of Theorem 2 inapplicable. Nonetheless, it is straightforward to check that the conditions of Theorem 1 still apply and, as a result, RPA is still guaranteed as long as the closed-loop system is stable. Furthermore, applying Eq. (2), by noting that *q* = [1 -10 0 0 1 2 1] ^*T*^, yields the setpoint expression given by *µ*_1_*/θ*_2_ (see Fig. 7(a)). Observe that the rate of expression *µ*_4_ of **Z**_**4**_ does not affect the setpoint — a result that is not immediate without resorting to Eq. (2). Similar to Fig. 4, the experimental results depicted in Fig. 7(b) demonstrate that the controller indeed ensures RPA yielding an average steady-state error of 3.9% over a wide dynamic range of setpoints compared to an error of 40.9% when operating in open loop.

**Fig. 7.**
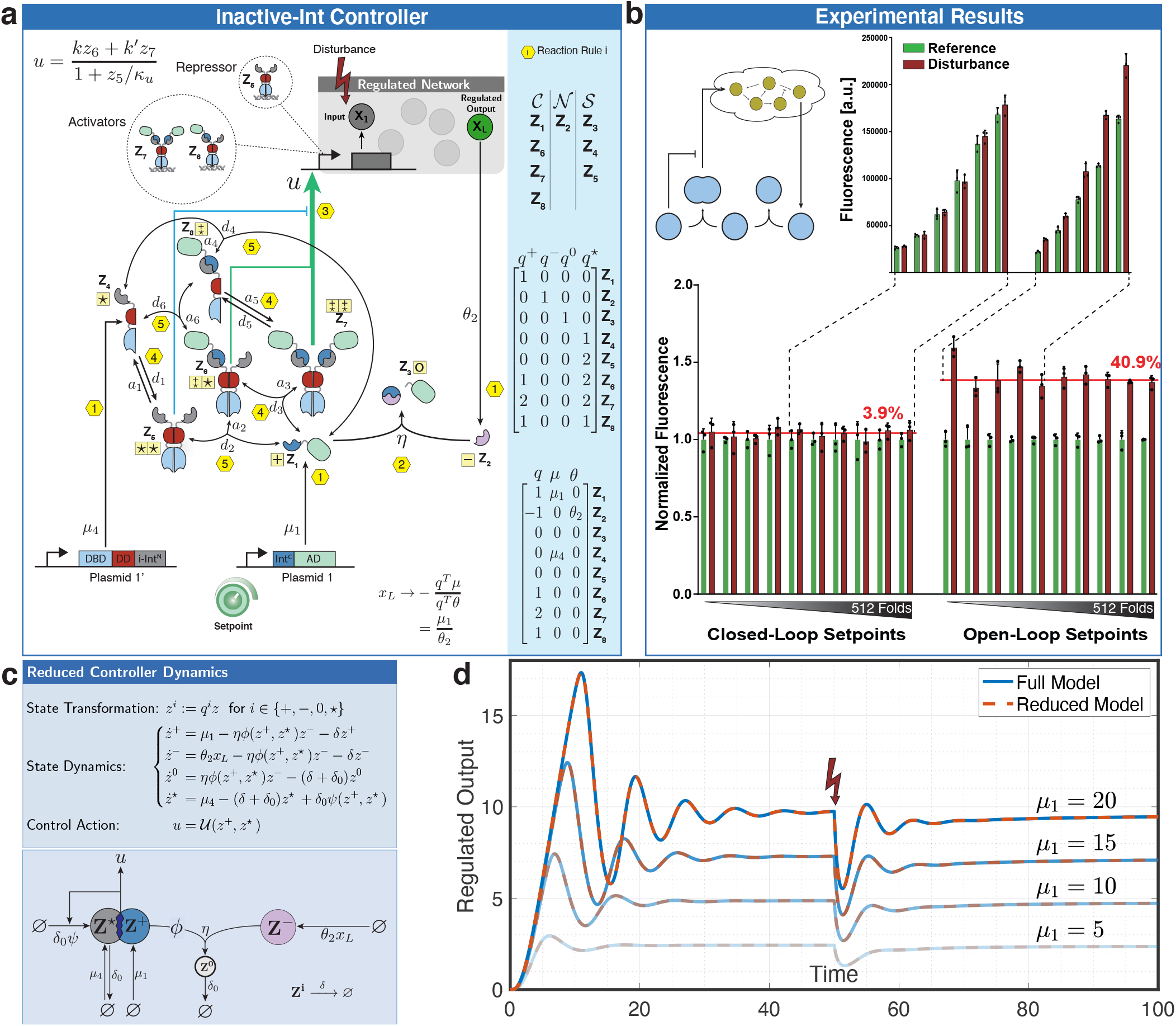
Inactive-intein controller: Theoretical and Experimental Analysis. (a) A schematic diagram of the inactive-intein controller. This controller consists of two genes, realized on separate plasmids. The gene in Plasmid 1 encodes for a protein (**Z**_**1**_) comprised of Int^C^-AD; whereas, the gene in Plasmid 1^*′*^ encodes for a protein (**Z**_**4**_) comprised of TetR-inactive Int^N^. Both genes are driven by a strong constitutive promoter (EF-1*α*), and their expression rates are denoted by *µ*_1_ and *µ*_4_, respectively. **Z**_**1**_ and **Z**_**4**_ can reversibly bind to form a heterodimeric transcription factor, which positively actuates the regulated network via the production of the input species **X**_**1**_. The production of the second split intein Int^N^, denoted by **Z**_**2**_, is driven by the regulated output **X**_**L**_ at a rate *θ*_2_ *x*_*L*_ to encode for the “sensing” reaction. When controller species contain a DD, reversible homo- or hetero-dimerization reactions occur with an association rate *a*_*i*_ and a dissociation rate *d*_*i*_. In this controller, only **Z**_**1**_ and **Z**_**2**_ can directly undergo the intein-splicing reaction, whose rate is denoted by *η*, because **Z**_**1**_ is the only species that contains an active Int^C^ segment not bound to the inactive Int^N^ segment. The spliced products **Z**_**3**_ do not have any activity. All controller species directly impacting the regulated network are indicated either as repressors or activators in the dashed bubbles, and the mathematical expression of the total control action *u* is shown as a (Hill-type) function of the repressors and activators. Every reaction is labeled from 1 to 6 according to the permitted reaction rules stated in Fig. 2. The entire charge matrix can be viewed in the blue shaded box where, additionally, all controller species have been grouped into the three classes, *C* -class, *N* -class and *S* -class, according to the species rules of Fig. 2. Since all the Species and Reaction Rules of Fig. 2 are respected, then by Theorem 1, we conclude that this controller ensures RPA (as long as closed-loop stability is maintained), and the setpoint can be shown, using Eq. (2), to be *µ*_1_ */θ*_2_ and is thus interestingly independent of *µ*_4_. (b) Steady-state error of the inactive-intein controller. The performance of the inactive-intein controller was tested in the closed-loop setup shown in Fig. 4(a). The only difference here is that the Int^N^ segment (Gp41-1) is replaced by an orthogonal Int^N^ (NrdJ-1) for the open-loop setting, and thus no intein-splicing reaction can occur. A simplified schematic (top left) of the controller topology is shown, and two bar graphs are sketched to report the experimental measurements in a fashion similar to that reported in Fig. 4. (c) Reduced Model. Unlike the three-dimensional reduced models in Fig. 6 that are obtained by directly applying Theorem 2, the reduced model here is four dimensional because it was necessary to introduce the dynamics of an additional state variable denoted by *z*^*⋆*^. The functions *ψ* and *ϕ* (not shown here) can be calculated by solving a set of algebraic equations. Note that the cartoon describing the reduced model is non-physical because the mathematical equations do not satisfy the structure of a simple motif like the models that satisfy Theorem 2. (d) Simulation Results. A closed-loop system is simulated for four increasing setpoints, where a model of a gene expression network is controlled by the inactive-intein controller. The simulations results demonstrate that the reduced model indeed accurately captures the dynamics of the original full model.

Although the model reduction recipe provided in Theorem 2 cannot be applied here, one can still invoke singular perturbation theory to this particular controller circuit to yield the reduced mathematical model depicted in Fig. 7(c). The model reduction here assumes, once again, that the reversible binding reactions are fast. Observe that, unlike the previous controllers, the reduced model is four dimensional. Intuitively, this is a result of an additional conservation law imposed by the inactive inteins which introduce an additional (fourth) vector *q*^*⋆*^ required to carry out the state transformation. Hence, the reduced mathematical model is described by the set of four ODEs for (*z*^+^, *z*^−^, *z*^0^, *z*^*⋆*^) shown in Fig. 7(c) where the functions *ϕ* and *ψ* can be calculated by solving a set of algebraic equations. A “fictitious network” describing the ODEs is also depicted in Fig. 7(c) to emphasize that the reduced model is mainly mathematical and cannot be easily translated to a simple motif. This highlights that controller circuits not adhering to the conditions of Theorem 2 fail to reduce to the simple motif given in Fig. 2. The reduced model is validated by the simulation results shown in Fig. 7(d) for four different setpoints and by applying a disturbance.

## Discussion

In this paper, we introduced a novel theoretical and experimental framework to design, build and analyze a broad class of biomolecular integral feedback controllers that achieve RPA. The framework is based on custom-built split inteins that are shown to be capable of realizing the sequestration reaction — the heart of the basic antithetic integral feedback motif — via protein splicing. The sequestration reaction in previously proposed (20, 37, 39, 40, 52) and built (6, 7, 9, 13) integral controllers, whether *in vivo*, or *in vitro*, relies on the complete stoichiometric annihilation of two controller species (see **Z**_**1**_ and **Z**_**2**_ in Fig. 1(a)). Here, we relax this requirement by establishing that the sequestration reaction does not have to fully annihilate the participating controller species, and, in fact, it suffices to stoichiometrically annihilate sub-components within these two controller species. Indeed, this is precisely what intein-splicing reactions do: active split inteins inserted in two target proteins are *inactivated* by undergoing the splicing reaction. While the function of the active split inteins is indeed *annihilated*, the spliced target proteins are still allowed to have specific functions. In fact, we showed that one can harness the function of the spliced proteins to augment the standalone integral controller with a filtered proportional component to yield a PI controller. We previously computationally demonstrated (see (38)) that the resulting filtered PI controller adds an extra degree of freedom which enables the enhancement of the transient performance while maintaining the RPA property. However, it is left for future work to back up this theory with experimental demonstrations. It is worth to mention that the realization of a molecular PI controller in mammalian cells is not new. Ideally, a proportional component can be theoretically achieved by appending the integral controller with an instantaneous negative feedback from the regulated output species **X**_**L**_ onto the input species **X**_**1**_ (see e.g. (37, 40)). This requires the output species **X**_**L**_ to have multiple functions including the production of **Z**_**2**_ for sensing and the inhibition of the input species **X**_**1**_ to realize the proportional component. In practice this might not be possible as the output species is determined by the biological application. In (7), this was circumvented by introducing additional genetic parts to express a proxy to the regulated output upon which the proportional control action is based on. Here, in contrast, the design flexibility and modularity offered by inteins allowed us to implement PI controllers by simply choosing an actuator and a suitable insertion site of the split-intein (see Fig. 3 and 6) without adding additional genetic parts and without requiring the regulated output **X**_**L**_ to have multiple functions.

The simple antithetic integral feedback control topology was first introduced in (20), and more recently a generalized antithetic topology was introduced in (6) which characterizes all RPA-achieving controllers involving exactly one sensing and one setpoint-encoding reaction. This characterization lead to simple algebraic conditions that enable RPA and are expressed in terms of quantities that are referred to as “charges”. The general charge analogy borrowed from electronics was made due to the lack of biological parts capable of respecting the algebraic conditions. This is exactly where inteins came in, because they naturally satisfy the RPA algebraic conditions and act as “charges” neutralizing each other via the intein-splicing reactions. In fact, Theorem 1, which is a direct application (tailored towards inteins) of a more general Theorem, is a generalization of the RPA sufficiency result of (6) such that multiple sensing and setpoint-encoding reactions are now allowed. Theorem 1 facilitates the screening of controller circuit designs for RPA. Furthermore, we went one step further here, beyond establishing RPA, to provide an easy-to-apply recipe for model reduction. The recipe is given in Theorem 2 which is, once again, a direct application (tailored towards inteins) of a more general Theorem (see Box.1 for an application example of these theorems in a purely mathematical and more general context, that is, without an intein-based interpretation). The model reduction result presented here exploits the time-scale separation imposed by fast reversible binding and conversion reactions and is established by invoking singular perturbation theory (54) together with the deficiency zero theorem (53) to prove structural stability of the slow manifold regardless of the rate parameters.

### Box 1.

Example Application of the RPA and Model Reduction Theorems to a General Controller Network

The goal of this illustrative example is to reduce the full model of the controller network comprised of *M* = 6 species **Z**_**1**_, **Z**_**2**_,…, **Z**_**6**_ to the simple motif comprised of **Z**^**+**^, **Z**^**–**^ and **Z**^**0**^ which is generally depicted in Fig.5. The full model here is chosen to be purely mathematical, i.e. with no relevance to inteins, in order to demonstrate that the theorems are not restricted to intein-based controllers only. Essentially, the *q* and (*q*^+^, *q*^−^, *q*^0^) vectors in Theorem 1 and 2 that were related to the number of inteins present in the controller species, are now more generally referred to as charge vectors inspired by (6). For establishing RPA, computing *q* := *q*^+^ *−q*^−^ is enough; whereas to carry out the model reduction technique, all individual positive, negative and neutral charge vectors (*q*^+^, *q*^−^, *q*^0^) need to be computed. Unlike the intein-based controller networks where the charge vectors are simply computed by counting intein segments, here we lay down general rules for constructing these charge vectors. Once these charge vectors are available, then one can easily apply Theorems 1 and 2 for establishing RPA and performing the model reduction, respectively. Note that in this example, we include three reversibly binding reactions but no conversion reactions for simplicity, and thus the conditions of Theorem 2 boil down to requiring the stoichiometry matrix of the binding network to be full-column rank and equal to *M−* 3 = 3. Furthermore, for simplicity, we start by assuming that **Z**_**1**_ and **Z**_**2**_ carries only one positive and negative charge, respectively. This assumption can however be relaxed to a more general number of charges.

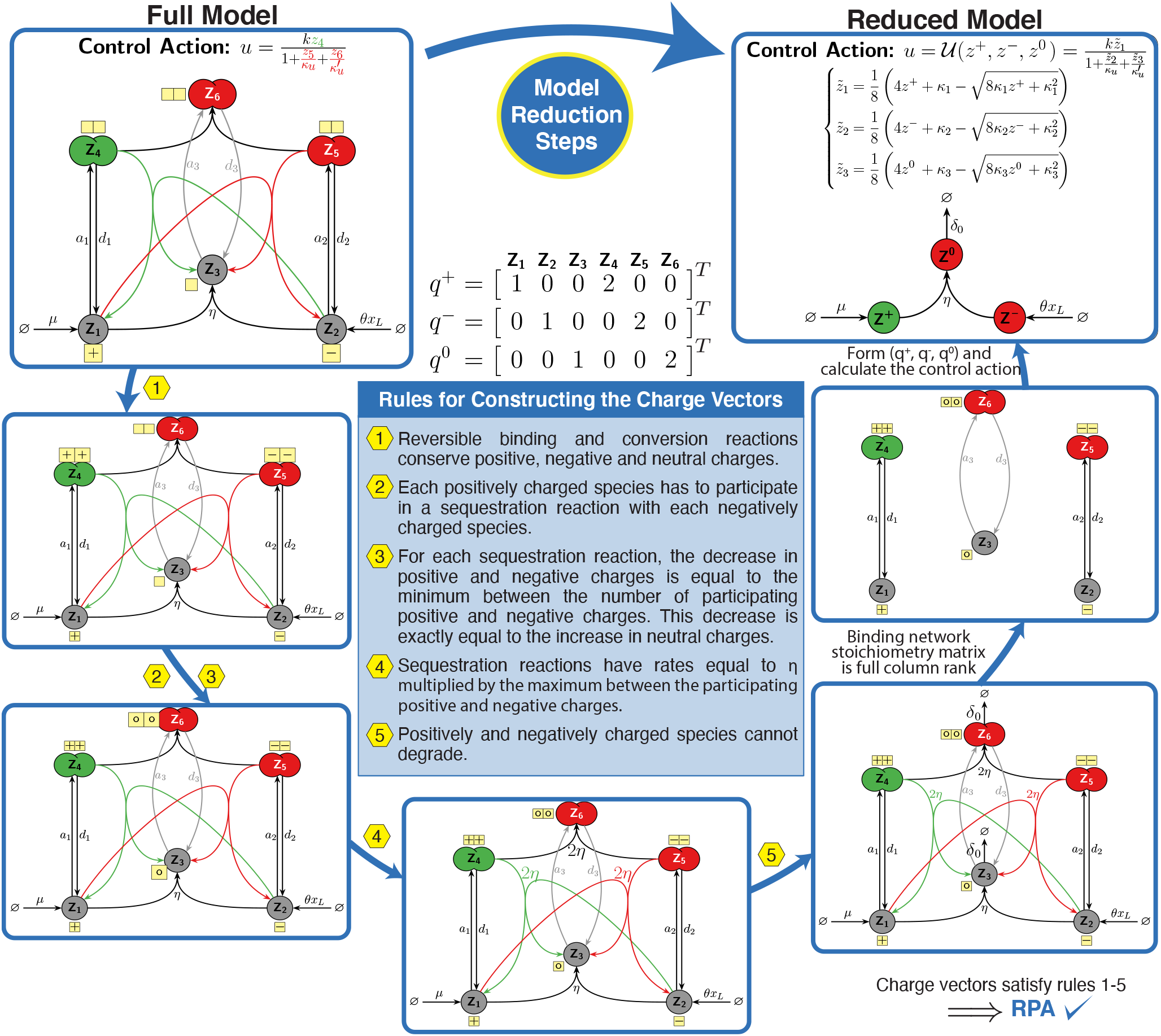

The five controller circuit implementations presented in this paper (see Fig. 3 and 7) are based on the widely used DNA binding domains TetR, ZF and Gal4. For the experimental verification of RPA, we used a simple regulated network (see Fig. 4(a)) that resulted in a two (resp. three) plasmid closedloop system depicted in Fig. 4 (resp. 7). The regulated network was intentionally chosen to be simple here, in order to minimize possible cross talks which might emerge from larger networks (e.g. burden)(48). This allowed us to focus our study on the controllers themselves instead of possible undesirable behaviors incurred by larger networks — an important topic that is not within the scope of the current study and is left for future work. Note that with this experimental setup, we were not able to directly detect the regulated output which is an mRNA (see Fig. 4(a)). To circumvent this, we used a fluorescent reporter which, unlike the regulated (mRNA) species, is not robust to translational burden.

The controller circuits that are designed, built and analyzed in this paper are all based on controller species generated using TFs. However, split inteins can also be introduced in other protein classes such as proteases and receptors. Split inteins can be even introduced in endogenous proteins to convert them into controller species. This has an attractive advantage of exploiting parts of the regulated network to realize the controller and, as a result, requiring less to no additional genes. From a protein engineering point of view, such designs may be more challenging than designs based on the well-characterized TFs used in this study. Besides tinkering with insertion sites, linker lengths and split-intein pairs, it is also possible to use more systematic approaches like transposon screens with inteins as performed by Ho et al. (56) or computationally-guided optimizations by Dolber et al. (57).

The remarkable flexibility offered by inteins for building integral controllers opens the doors to many possible future research directions. For instance, it is easy to think of regulated networks with negative gain, in other words, producing more input species **X**_**1**_ leads to a lower concentration of the regulated output species **X**_**L**_. For example, producing more insulin leads to a lower concentration of glucose in the blood. As a result, to realize an overall negative feedback, the actuation direction of the controller species **Z**_**1**_ would have to be flipped, that is instead of having **Z**_**1**_ upregulating **X**_**1**_ (like in Fig. 1(a) and, in fact, all previously built antithetic integral controllers), **Z**_**1**_ would have to downregulate **X**_**1**_. Intein-based realizations of such “negative actuation” mechanisms can be easily carried out using repressors or proteases.

Another possible future direction is intein-based implementations of more advanced controllers. For example, one can easily add functional domains to the controller species **Z**_**2**_, which was comprised of a standalone Int^N^ segment in all the controller circuits proposed here. These added domains enable the implementation of the rein controller introduced in (58) which is capable of enhancing the overall performance. Another example is the implementations of more advanced biomolecular Proportional-Integral-Derivative (PID) controllers (37) that are capable of shaping the transient response and reducing cell-to-cell variability. In particular, the wide library of orthogonal split inteins (50) allows one to implement the fourth order PID controller (37) that is comprised of two antithetic motifs: antithetic integrator and antithetic differentiator.

In conclusion, rather than providing another way of implementing antithetic integral controllers, we propose here a novel systematic (theoretical and experimental) approach of designing, building and analyzing a broad class of biomolecular integral controllers that are capable of achieving RPA. The key of our approach is the exploitation of the splicing reactions that occur between split inteins. Due to their simplicity, modularity, irreversibility, lack of side effects and applicability across species, we believe that inteins will revolutionize biomolecular controllers and partake in filling the gap between theory and experiments.

## Materials and Methods

### Plasmid Construction

All plasmids were generated with a mammalian adaptation of the modular cloning (MoClo) yeast toolkit standard (59). All individual parts were generated by PCR amplification (Phusion Flash High-Fidelity PCR Master Mix; Thermo Scientific) or synthesized with Twist Bioscience. The parts were then assembled with golden gate assembly. All enzymes for plasmid construction were obtained from New England Biolabs (NEB). Constructs were chemically transformed into *E. coli* Top10 strains.

### Cell culture

All experiments were performed with HEK293T cells (ATCC, strain number CRL-3216). The cells were cultured in Dulbecco’s modified Eagle’s medium (DMEM; Gibco) supplemented with 10 % FBS (Sigma-Aldrich), 1x GlutaMAX (Gibco), 1 mm Sodium Pyruvate (Gibco), penicillin (100U/*µ*L), and streptomycin (100 *µ*g/mL) (Gibco) at 37° with 5 % CO2. The cell culture was passaged into a fresh T25 flask (Axon Lab) every 2 to 3 days. Upon detachment some part of the cell suspension was used for the transfection.

### Transfection

All plasmids were isolated using ZR Plasmid MiniprepClassic (Zymo Research). The plasmids were introduced to the HEK293T cells via suspension transfection. A transfection solution in Opti-MEM I (Gibco) was prepared using Polyethylenimine (PEI) “MAX” (MW 40000; Polysciences, Inc.) at a 1:3 (*µ*g DNA to *µ*g PEI) ratio while the culture was detached with Trypsin-EDTA (Gibco). The cell density was assessed with the automated cell counter Countess II FL (Invitrogen). 100 *µ*L of culture with 26’000 cells was transferred in each well of the plate Nunc Edge 96-well plate (Thermo Scientific). The transfection mixture was added to the cells once it has incubated for approximately 30 min.

### Flow cytometry

The cells were detached approximately 48 hours after transfection on the Eppendorf ThermoMixer C at 25°C at 700 rpm with 53 *µ*L Accutase solution (Sigma-Aldrich) per well for 20 min. The fluorescence data was collected on the Beckman Coulter CytoFLEX S flow cytometer with the 488 nm excitation with a 525/40+OD1 bandpass filter and the 638 nm excitation with a 660/10 bandpass filter. All data was processed with the CytExpert 2.3 software.

## ACKNOWLEDGMENTS

This project has received funding from the European Research Council (ERC) under the European Union’s Horizon 2020 research and innovation programme (CyberGenetics; grant agreement 743269). The authors would like to thank Dr. Zhou Fang for the insightful discussions.

## COMPETING INTERESTS

ETH Zurich has has filed a patent application on behalf of the inventors S.A., M.F., C.H.C. and M.K. on the genetic circuit designs described (application no. EP22186956.3).

